# Retinotopic Organization of Scene Areas in Macaque Inferior Temporal Cortex

**DOI:** 10.1101/131409

**Authors:** Michael J. Arcaro, Margaret S. Livingstone

## Abstract

Primates have specialized domains in inferior temporal (IT) cortex that are responsive to particular image categories. Though IT traditionally has been regarded as lacking retinotopy, several recent studies in monkeys have shown that retinotopic maps extend to face patches along the lower bank of the superior temporal sulcus (STS) in IT cortex. Here, we confirm the presence of visual field maps within and around the lower bank of the STS and extend these prior findings to scene-selective cortex in the ventral-most regions of IT. Within the occipito-temporal sulcus (OTS), we identified two retinotopic areas, OTS1 and OTS2. The polar angle representation of OTS2 was a mirror reversal of the OTS1 representation. These regions contained representations of the contralateral periphery and were selectively active for scene vs. face, body, or object images. The extent of this retinotopy parallels that in humans and shows that the organization of the scene network is preserved across primate species. In addition retinotopic maps were identified in dorsal extrastriate, posterior parietal, and frontal cortex as well as the thalamus, including both the LGN and pulvinar. Taken together, it appears that most, if not all, of the macaque visual system contains organized representations of visual space.

**SIGNIFICANCE STATEMENT:** Primates have specialized domains in inferior temporal (IT) cortex that are responsive to particular image categories. Though retinotopic maps are considered a fundamental organizing principle of posterior visual cortex, IT traditionally has been regarded as lacking retinotopy. Recent imaging studies have demonstrated the presence of several visual field maps within lateral IT. Using neuroimaging, we found multiple representations of visual space within ventral IT cortex of macaques that included scene-selective IT cortex. The scene domains were biased towards the peripheral visual field. These data demonstrate the prevalence of visual field maps throughout the primate visual system, including late stages in the ventral visual hierarchy, and support the idea that domains representing different categories are biased towards different parts of the visual field.

## INTRODUCTION

In the primate brain, object categories are represented in a large swath of IT within and around the superior temporal sulcus (STS). Macaque IT cortex contains specialized domains that are responsive to particular biologically important categories such as faces(Tsao et al., 2003; Rajimehr et al., 2009), bodies(Pinsk et al., 2005; Bell et al., 2009; Pinsk et al., 2009), or places(Nasr et al., 2011; Kornblith et al., 2013). This part of cortex had been generally regarded as lacking any other large-scale functional organization. However, recent fMRI studies have shown that visual field maps exist within and around the lower bank of the STS(Janssens et al., 2014; Kolster et al., 2014) as well as within ventral parts of IT(Orban et al., 2014). Some of these maps partially overlap with face-selective domains(Janssens et al., 2014). Here, we investigated the extent of this retinotopy in ventral parts of IT and its relationship to scene-selective domains.

In humans, several visual field maps have been identified within the ventral-most regions of category-selective temporal cortex. Anterior to human V4 (hV4), in ventral occipital cortex, two visual field maps, VO1 and VO2, have been identified that have similar response selectivity for color and object-related information(Brewer et al., 2005; Wandell et al., 2005). Anterior to these two maps within the collateral sulcus, two additional maps, PHC1 and PHC2, have been identified that mainly represent peripheral space and are selective for scenes vs. other categories(Arcaro et al., 2009). Many parallels in cortical organization have been proposed between humans and monkeys(Van Essen et al., 2001; Orban et al., 2004; Arcaro and Kastner, 2015; Orban, 2016). We further evaluate the similarity of visual cortical organization between primate species by comparing the functional organization of medial ventral temporal.

Here, we performed retinotopic mapping in four macaque monkeys. In addition to previously identified maps, we identified two new visual field maps, which we will refer to as OTS1 and OTS2. These new regions were located ventral and medial to retinotopic areas V4A and PIT. Both regions contained representations of the entire contralateral periphery. OTS1 and OTS2 were in the same location as the functionally defined scene area, LPP(Kornblith et al., 2013) and were selectively responsive to scenes compared to objects, faces, or bodies. Further, the visual field organization of OTS1/2 corresponded to the organization of scene-selective retinotopic areas PHC1/2 in humans(Arcaro et al., 2009). Our data provide novel evidence that monkey LPP is the homologue to human area PPA, and demonstrate that the retinotopic organization of macaque IT cortex is more extensive than previously described.

## MATERIALS AND METHODS

### Monkeys

Functional MRI studies were carried out on 4 juvenile Macaca mulattas, 2 female and 2 male, born in our laboratory. All procedures were approved by the Harvard Medical School Animal Care and Use Committee and conformed with National Institutes of Health guidelines for the humane care and use of laboratory animals. Three monkeys were co-housed with their mothers in a room with other monkeys for the first 4 months, then co-housed with other juveniles, also in a room with other monkeys. As part of a separate experiment, the other monkey (M3) was hand reared by humans for the first year, then was co-housed with other juveniles. For scanning they were alert, and their heads were immobilized using a foam-padded helmet with a chinstrap that delivered juice. The monkeys were scanned in a primate chair that allowed them to move their bodies and limbs freely, but their heads were restrained in a forward-looking position by the padded helmet. The monkeys were rewarded with juice for maintaining a central fixation within a 2° window. Gaze direction was monitored using an infrared eye tracker (ISCAN, Burlington, MA).

### Stimuli

Visual stimuli were projected onto a screen at the end of the scanner bore.

#### Retintopic Mapping

Retinotopic mapping was performed in all monkeys > 1.5 years of age when they were able to maintain fixation for extended periods of time.

To obtain polar angle maps, visual stimuli consisted of a wedge that rotated either clockwise or counterclockwise around a central fixation point. The wedge spanned 0.5-10° in eccentricity with an arc length of 45° and moved at a rate of 9°/s. The wedge consisted of a colored checkerboard with each check’s chromaticity and luminance alternating at the flicker frequency of 4 Hz (See Arcaro et al., 2011 for details). Each run consisted of eight cycles of 40 s each. 10-12 runs were collected with an equal split in the direction of rotation.

To obtain eccentricity maps, visual stimuli consisted of an annulus that either expanded or contracted around a central fixation point. The duty cycle of the annulus was 10%; that is, any given point on the screen was covered by the annulus for only 10% of the time. The annulus swept through the visual field linearly. The ring consisted of the same colored checkerboard as the wedge stimulus. Each run consisted of seven cycles of 40 s each with 10 s of blank, black backgrounds in between. Additional blank periods were inserted to distinguish responses between the foveal and most peripheral positions. 10-12 runs were collected with an equal split in direction.

#### Static Images

Responses to image categories of scenes, faces, bodies, and inanimate objects were probed. Each scan comprised blocks of each image category; each image subtended 20°×20° of visual angle and was presented for 0.5 seconds; block length was 20 seconds, with 20 seconds of a neutral gray screen between image blocks. Blocks and images were presented in a counterbalanced order. The scene images were of familiar laboratory scenes, the faces were mosaics of monkey faces on a pink-noise background, with faces covering most of the 20 degree image; bodies were mosaics of headless monkey bodies on pink noise, and objects were mosaics of familiar objects on pink noise, All images were equated for spatial frequency and luminance using the SHINE toolbox(Willenbockel et al., 2010).

### Scanning

Monkeys were scanned in a 3-T TimTrio scanner with an AC88 gradient insert using 4-channel surface coils (custom made by Azma Maryam at the Martinos Imaging Center). Each scan session consisted of 10 or more functional scans. We used a repetition time (TR) of 2 seconds, echo time (TE) of 13ms, flip angle of 72°, iPAT = 2, 1mm isotropic voxels, matrix size 96x96mm, 67 contiguous sagittal slices. To enhance contrast(Vanduffel et al., 2001; Leite et al., 2002), we injected 12 mg/kg monocrystalline iron oxide nanoparticles (Feraheme, AMAG Parmaceuticals, Cambridge, MA) in the saphenous vein just before scanning.

### General preprocessing

Functional scan data were analyzed using Analysis of Functional NeuroImages (AFNI; RRID:nif-0000-00259)(Cox, 1996), SUMA(Saad and Reynolds, 2012), Freesurfer (Freesurfer; RRID:nif-0000-00304)(Dale et al., 1999; Fischl et al., 1999), JIP Analysis Toolkit (written by Joseph Mandeville), and MATLAB (Mathworks, RRID:nlx_153890). Each scan session for each monkey was analyzed separately. All images from each scan session were AFNI-aligned to a single timepoint for that session, detrended and motion corrected. Data were spatially filtered using a Gaussian filter of 2 mm full-width at half-maximum (FWHM) to increase the signal-to-noise ratio (SNR) while preserving spatial specificity. Each scan was normalized to its mean. Data were registered using a two-step linear then non-linear alignment approach (JIP analysis toolkit) to a high-resolution (0.5mm) anatomical image acquired for monkeys M1 and M2 (> 1.5 years) and to a standard anatomical template (F99) for monkeys M3 and M4. First, a 12-parameter linear registration was performed between the mean RPI image for a given session and a high-resolution anatomical image. Next, a nonlinear, diffeomorphic registration was conducted. To improve registration accuracy of ventral cortex, we manually drew masks that excluded the cerebellum for both EPIs and anatomicals prior to registration.

### Retinotopy Analysis

Fourier analysis was used to identify spatially selective voxels from the polar angle and eccentricity stimuli(Bandettini et al., 1993; Engel et al., 1997). For each voxel, the amplitude and phase (the temporal delay relative to the stimulus onset) of the harmonic at the stimulus frequency were determined by a Fourier transform of the mean time series. To correctly match the phase delay of the time series of each voxel to the phase of the wedge/ring stimuli, and thereby localize, the region of the visual field to which the underlying neurons responded best, the response phases were corrected for a hemodynamic lag (4 s). The counterclockwise (expanding) runs were then reversed to match the clockwise (contracting) runs and averaged together for each voxel.

An F-ratio was calculated by comparing the power of the complex signal at the stimulus frequency to the power of the noise (the power of the complex signal at all other frequencies). Statistical maps were threshold at a *p* < 0.0001 (uncorrected for multiple comparisons). Data were registered and projected onto individual surfaces for monkeys M1 and M2. Monkeys M3 and M4 did not have individual surfaces, so data were projected onto a standard surface template (F99, (Van Essen and Dierker, 2007)). When displaying phase estimates, a 20-point color scale was assigned to the polar angle datasets with each color representing 18° visual angle, and a 20-point color scale was assigned to the eccentricity datasets with each color representing 1.6° eccentricity. Contiguous clusters of spatially selective voxels were identified throughout cortex. Borders between visual field maps were identified based on reversals in the systematic representation of visual space, particularly with respect to polar angle. Eccentricity representations were evaluated to ensure that phase progressions were essentially orthogonal (nonparallel) to the polar angle phase progression and to differentiate between the MT+ cluster(Kolster et al., 2009) and surrounding extrastriate visual field maps. Activations were found in both dorsal and ventral striate and extrastriate cortex. Though dorsal and posterior regions were outside the focus of our analysis, to validate our mapping, a series of visual field maps (Fig. 1) was identified in general accordance with previous literature(Kolster et al., 2009; Arcaro et al., 2011; Janssens et al., 2014). Surface area was estimated as the average surface area for pial and white matter surfaces segmentations using AFNI’s SurfMeasures. Surface estimates were derived only for monkeys M1 and M2, and monkeys M3 and M4 were projected onto a standard template surface.

**Figure 1.**
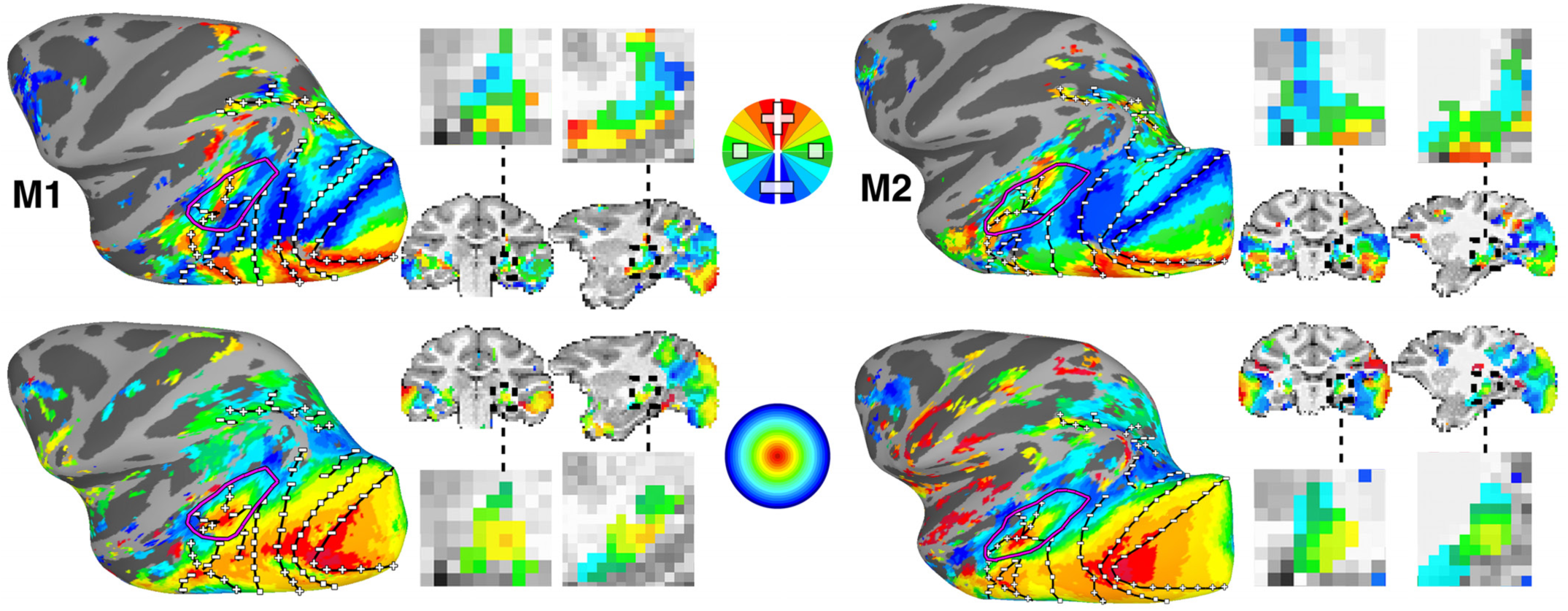
Retinotopic maps across occipital, temporal, and parietal cortex as well as the thalamus. (top) Polar angle and (bottom) eccentricity maps are shown on inflated surface views of lateral cortex in the left hemispheres and sagittal and coronal volume slices of monkeys M1 and M2. Full volume slices are accompanied with zoomed in views of the ventral thalamus. The color code indicates the phase of the fMRI response and therefore the preferred region of the visual field. For polar angle maps, only contralateral representations are displayed. Data were threshold at p < 0.0001. Black solid lines mark meridian borders between visual field maps. Black dashed line in volume image highlights a region of the posterior thalamus that includes the LGN and pulvinar. Plus, minus, and square symbols mark upper, lower and horizontal meridians. The purple line encompasses the MT cluster (comprising MT, MST, FST, and LST)(Kolster et al., 2009).

To compute the representation of the visual field as a function of polar angle in our areas of interest, the visual field was divided into four sectors: contralateral vs. ipsilateral, and upper vs. lower. To compute the representation of the visual field as a function of eccentricity, the visual field was divided into three equally spaced sectors spanning foveal (0-3.33°), midfield (3.34-6.66°), and peripheral (6.67-10°) visual space. The number of voxels within each sector was tallied and divided by the total number of voxels in each area of interest to derive the percentage visual field coverage for each subject. Data were collapsed across hemispheres and averaged across monkeys to derive a group mean average. Comparisons were made between visual field sectors and two-tailed, t-tests were used to assess statistical significance.

To directly compare visual field maps across monkeys, each monkey’s data was aligned to a standard template (F99(Van Essen and Dierker, 2007)) surface using nonlinear registration (JIP Analysis Toolkit). To derive group average polar angle and eccentricity maps, individual polar angle and eccentricity visual field maps were converted to Cartesian space, averaged, and threshold such that all voxel’s were required to be significant in 3/4 monkeys. Group average maps were compared with the borders of the Lewis and Van Essen macaque F99 atlas(Lewis and Van Essen, 2000; Van Essen and Dierker, 2007).

### Stimulus Category Analysis

A multiple regression analysis (AFNI’s 3dDeconvolve (Cox, 1996)) in the framework of a general linear model (Friston et al., 1995) was performed on the category experiments for each monkey separately. Each stimulus condition was modeled with a MION-based hemodynamic response function(Leite et al., 2002). Additional regressors that accounted for variance due to baseline shifts between time series, linear drifts, and head motion parameter estimates were also included in the regression model. Due to the time-course normalization, beta coefficients were scaled to reflect percent signal change. Since MION inverts the signal, the sign of beta values were inverted to follow normal fMRI conventions of increased activity are represented by positive values. Brain regions that responded more strongly to scenes, monkey faces, monkey headless bodies or familiar objects were identified by contrasting presentation blocks of each of these image categories. Maps of beta coefficients were clustered (>10 adjacent voxels) and threshold at p<0.00001 (FDR-corrected).

D-primes were calculated from beta-coefficients. Critical t values were used to calculate standard errors. These values were used to calculate a d-prime index using the following d’ formula (Afraz et al., 2006; Grill-Spector et al., 2006):

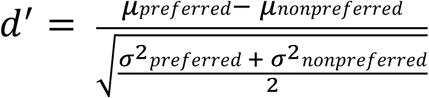

where *μpreferred* and *μnonpreferred* are -1 (to invert the MION response) x the average responses to the preferred stimulus category (e.g., scenes) and the average responses to the nonpreferred stimulus category (e.g., objects); *σpreferred* and *σnonpreferred* are the standard deviations.

## RESULTS

Polar angle and eccentricity maps were measured for the central 10° of the visual field using a smoothly rotating wedge stimulus and an expanding annulus stimulus, in monkeys trained to maintain fixation. Bilateral activations within occipital, temporal, and parietal cortex were found in both hemispheres of all four monkeys for both polar angle and eccentricity mapping. Individual activation maps of polar angle and eccentricity are shown overlaid on lateral views of inflated surface reconstructions for the left (LH) hemispheres in two example monkeys (Fig. 1). For the polar angle mapping, activations within each hemisphere were mainly confined to the contralateral hemifield. The color of each voxel was determined by the phase of its responses and indicates the region of the visual field to which that voxel was most responsive. For polar angle maps, the upper visual field (UVF) is denoted in red-yellow, the horizontal meridian (HM) in green, and the lower visual field (LVF) in light and dark blue. For eccentricity maps, the central space (0-3.5°) is denoted in red/orange, mid-eccentricities in green (3.5-6°), and the periphery in blue (6-10°).

We identified a series of visual field maps throughout occipital, temporal, and parietal cortex in each monkey, consistent with previous fMRI mapping studies in monkeys(Brewer et al., 2002; Kolster et al., 2009; Arcaro et al., 2011; Janssens et al., 2014; Kolster et al., 2014) (Fig. 1). Additional representations of contralateral visual space were identified within the intraparietal sulcus, frontal cortex, and the thalamus in all monkeys. Representations of the upper and lower vertical meridians were identified anterior to the previously reported LIP visual field map(Patel et al., 2010; Arcaro et al., 2011), suggesting the presence of at least two additional visual maps within the inferior bank of the IPS. Consistent with a recent fMRI study(Janssens et al., 2014), contralateral representations of visual space were also identified within the ramus of the arcuate corresponding to the frontal eye fields area. Contralateral representations of visual space were also identified in both the lateral geniculate nucleus (LGN) and ventral pulvinar of the posterior thalamus(Kaas et al., 1972; Malpeli and Baker, 1975; Bender, 1981). Both thalamic regions showed an inversion of the visual field with the UVF located ventrally and the LVF located dorsally. Within the pulvinar, a foveal representation was found in the middle of the extent of the polar angle maps with increasingly peripheral representations extending both laterally and medially. This is consistent with the proposal of two visual field maps within the ventral pulvinar(Bender, 1981; Shipp, 2003). The retinotopic organization of the LGN and pulvinar in our macaques was also consistent with prior imaging studies in humans(Schneider et al., 2004; Arcaro et al., 2015; DeSimone et al., 2015), confirming that the organization of the visual thalamus is similar across primate species. Retinotopically-specific activity almost completely covered visually-evoked activity (identified from the category stimulus experiment) with the exception of anterior temporal STS (AIT), which was visually activated, but did not show clear retinotopic responses. Overall, visual field maps were prevalent throughout the extent of the visual system, underscoring the importance of topographic representations as an organizing principle for sensory systems.

### Ventral V1, V2, V3, V4, V4A, and PIT

For the purposes of this study, we focused on activations within medial ventral temporal cortex (Fig. 2). Ventral and medial to areas V4, V4A and PITv, we identified two visual field maps within the occipitotemporal sulcus (OTS), which we refer to as OTS1 and OTS2. Below, we provide a brief description of the retinotopic organization surrounding OTS1/2 and a more detailed description of these two new areas.

**Figure 2.**
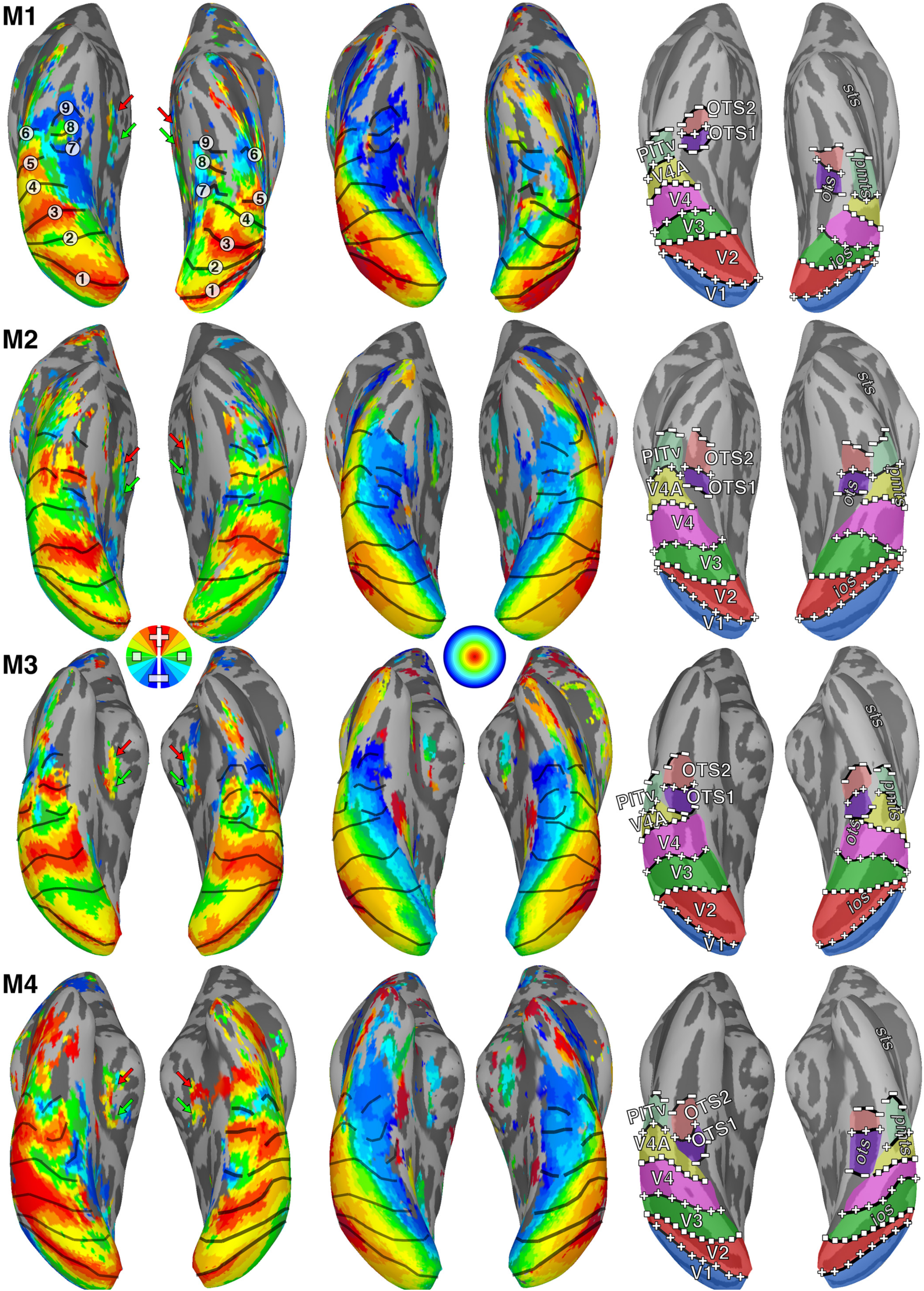
Retinotopic maps in ventral temporal cortex of four monkeys. (left) Polar angle and (middle) eccentricity maps are shown in inflated surface views of ventral occipital and temporal cortex. The color code indicates the phase of the fMRI response and therefore the preferred region of the visual field. For polar angle maps, only contralateral representations are displayed. Data were threshold at p < 0.0001. Black solid lines mark the borders between visual field maps. In M1, the locations of 6 reversals in polar angle progression corresponding to areal borders are labeled. (right) Areal extent for newly defined areas OTS1 and OTS2 as well as the ventral halves of V1, V2, V3, V4, V4A and the entire area of PITv are shown. Plus, minus, and square symbols mark upper, lower and horizontal meridians. Red and green arrows in the polar angle maps indicate partial surface projections of LGN and pulvinar visual maps, respectively.

In agreement with previous reports(Brewer et al., 2002; Fize et al., 2003; Arcaro et al., 2011), visual areas V1, V2, V3, and V4 were identified within occipital cortex in each hemisphere for all four monkeys. The UVF and LVF representations were noncontiguous, split between the ventral and dorsal cortex, respectively (Figs. 1 & 2). Within ventral occipital and temporal cortex, a clear alternating progression of visual field was identified along the anterior-posterior axis. Areas were differentiated based on reversals in visual field progression with respect to the polar angle axis that were orthogonal to the eccentricity axis(Sereno et al., 1995). Altogether, five reversals in polar angle phase progression were identified as the vertical and horizontal meridian borders (VM & HM) between visual areas (see M1 in Fig. 2, left column). Starting from within the calcarine sulcus, the HM of the V1 visual map progressed anterior to (#1) an upper vertical meridian that forms the border with area V2 then reverses back to (#2) an HM that corresponds to the border between V2 and V3, and then reverses again (#3) to another upper vertical meridian that forms the border between V3 and V4. Anterior to V4, the visual field continues along lateral parts of occipital cortex into the posterior STS, reversing back to (#4) an HM, forming the border between V4 and V4A, and then reversing again to (#5) an upper vertical meridian that forms the border between V4A and PITv. The visual field representation reverses once more and progresses all the way to (#6) a lower vertical meridian, forming the anterior border of PITv; which comprises a full hemifield representation of the UVF and LVF in contrast to the UVF-only quarter-field representations of V1-V4A. As previously described, PITv was located within the posterior medial temporal sulcus (PMTS) in all four monkeys(Janssens et al., 2014; Kolster et al., 2014).

The progression of eccentricity representations within ventral occipital and temporal cortex (Fig. 1, bottom row & Fig. 2, middle column) was orthogonal to the polar angle map. There was a large swath of central visual field representation (red/yellow), extending anteriorly along the lateral surface from V1 in occipital cortex to the lower bank of the STS in temporal cortex. From the representation of central space, eccentricity representations progressed ventral and medial to mid-eccentricities and then back to the periphery. Two peaks in the foveal representations (red) were consistently identified (Fig. 1, middle column); one where the dorsal and ventral halves of V1-V4 converged (meridians indicated as 1-4) and the other where PITv (meridians 5-6) converged with areas PITd and OTd as previously reported(Janssens et al., 2014; Kolster et al., 2014). Dorsal and ventral portions of area V4A (meridians 4-5) converged in-between these two foveal peaks. Anterior to these two foveal peaks, an additional foveal representation was identified along the lower bank of the STS in all monkeys.

#### Areas OTS1 and OTS

### Polar angle and eccentricity maps in the occipitotemporal sulcus

Medial to the peripheral representations of V4 and V4A, two representations of contralateral visual space were found within the OTS of each hemisphere, which we refer to as OTS1 and OTS2 (Fig. 2). A LVF representation medial to areas V4 and V4A (Fig. 2; #7 in M1) was identified as the posterior border of OTS1 in all monkeys. In 6 of 8 hemispheres, a representation of the lower vertical meridian (dark blue) was identified. In the right hemisphere of monkey M3, the posterior part of OTS1 mainly contained HM representations and the posterior border was defined at the HM representation that extended furthest into the lower quadrant of the visual field^1^. OTS1 was defined by a polar angle progression extending from this LVF representation anterior to (#8) an UVF representation that marked the border between OTS1 and OTS2. In 4 of 8 hemispheres, a representation of the upper vertical meridian (red) was identified. OTS2 was defined by polar angle phase progressions extending from the midpoint of the UVF representation shared by OTS1 to (#9) a LVF representation further anterior. In 8 of 8 hemispheres, a representation of the LVF vertical meridian (dark blue) was identified. In the right hemisphere of M2 and left hemisphere of M3, the representation of visual space in OTS2 was sparse at this threshold, but the general organization was consistent with the other hemispheres/monkeys. In all monkeys, OTS1 and OTS2 contained almost exclusively peripheral representations (> 7°) and there was no clear progression of visual field representation along the eccentricity dimension. Medial to OTS1/2 there was a small patch of cortex lacking significant visual field representations, then a jump back to a foveal representation in 5 out of 8 hemispheres. Since we included a 10s gap between presentations of foveal and peripheral ring stimuli, it is unlikely that this foveal representation is an artifact of "wrap-around" that can affect phase-encoded paradigms. There was no clear polar angle organization or progression to midfield/peripheral representations within this part of medial cortex and it thus appeared to be distinct from the OTS maps. Alternatively, this foveal representation may be part of the OTS maps; due to the large receptive fields and relatively small cortical size, we may not have had the sensitivity to resolve polar angle organization in this region.

To further investigate the topographic organization within OTS1 and OTS2, the polar angle phase progression was quantified. Typically for such an analysis, linear ROIs are drawn either along iso-polar angle lines orthogonal to eccentricity(Arcaro et al., 2009; Arcaro et al., 2011) or along iso-eccentricity lines orthogonal to polar angle(Schluppeck et al., 2005; Silver et al., 2005; Kolster et al., 2009; Kolster et al., 2014). In both cases, polar angle phase values are plotted as a function of distance from the starting point. Since there was no clear progression of eccentricity organization within OTS1 and OTS2, line segments were drawn orthogonal to the polar angle axis along the lateral (red dots) and medial borders (blue dots) as well as along the midline (green dots) in both maps as indicated by the schematic lines in Figure 3a. The polar angle phases along these line ROIs were plotted as a function of cortical distance from the posterior border of OTS1. In the Figure 3a plot, dots correspond to individual nodes located along each line segment in monkey M4 as indicated in the schematic above and the black line corresponds to the average across the three line segments. The polar angle representations started within the LVF (blue) and progress to the UVF (red) corresponding to the border between OTS1 and OTS2, then reversed back to the LVF (Fig. 3a, bottom). This analysis was performed for each hemisphere in each monkey separately. The polar angle representations along each ROI were strongly correlated within individual hemispheres (mean r = 0.83), demonstrating a consistency in the visual field organization within individual maps. Polar angle phase values between area borders were then interpolated into a common space, which allowed for group averaging (Fig. 3b). For the group average plots (Fig. 3b), each colored line corresponds to an individual monkey (hemispheres plotted separately) and the black line corresponds to the group average. It is evident in both the individual monkeys and group average that the posterior border of OTS1 and anterior border of OTS2 corresponded to troughs in the plot (closest to the LVF meridian) and the shared border between OTS1 and OTS2 corresponded to the peak of the plot (closest to the UVF meridian). Even in monkey M1 (red line) that had mainly LVF representations (Fig. 2), the polar angle map progressed from a LVF representation at the posterior border of OTS1 to UVF representations at the identified border or OTS1 and OTS2, and then reversed back to a LVF representation at the border of OTS2. Individual monkey polar angle phase progressions were strongly correlated (r = 0.90), demonstrating a consistency in the visual field organization of OTS1 and OTS 2 across monkeys.

**Figure 3.**
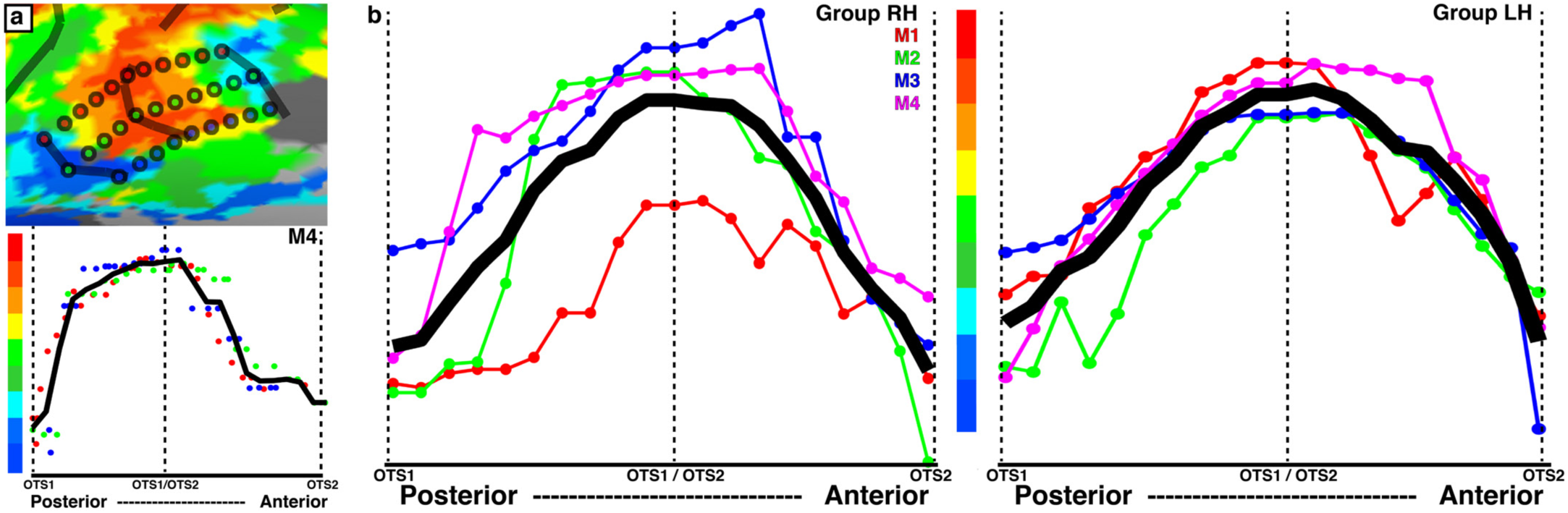
Analysis of topographic organization within OTS1 and OTS2. (a) Polar angle maps for OTS1 and OTS2 in the RH of monkey M4. (a, top) Red, green, and blue dotted lines correspond to the ROIs along the lateral, midline, and medial borders of OTS1 and OTS2. (a, bottom) Polar angle phase values plotted as a function of cortical distance (relative to posterior border of OTS1) for all three linear ROIs and the average (black line). (b) Polar angle phase plot for the RH and LH hemispheres of individual monkeys (averaged across all 3 lines within each hemisphere as illustrated in a) and the group average (black line). Phase values were interpolated into a common space, which allowed for averaging across monkeys. The smooth progression of phase values within the OTS maps and the phase reversals at the area boundaries were apparent in the group average as well as in the individual subjects.

The anatomical extent of OTS1 and 2 was consistent between hemispheres and monkeys as is shown on the surface (Fig. 2) as well as within the volume (Fig. 4). On average, the borders of OTS1 extended from 5.5 to -1.0 (A-P), 1.75 to 7.5 (I-S) and 17.0 to 23.0 (M-L), and for OTS2 from 9.5 to 3.5 (A-P), 0.5 to 6.0 (I-S) and 17.0 to 23.0 (M-L). The mean surface area estimates (average between pial and white matter surface segmentations) for OTS1 and OTS2 were 45.1 +/- 3.8 mm^2^ and 40.3 +/- 3.1 mm^2^, respectively, which corresponded to ∼4% of V1’s surface area. The mean surface volume estimates (measured between pial and white matter) for OTS1 and OTS2 were 58.6 +/- 4.2 mm^3^ and 66.6 +/- 3.7 mm^3^, respectively, which corresponded to between 6-7% of V1’s surface volume.

**Figure 4.**
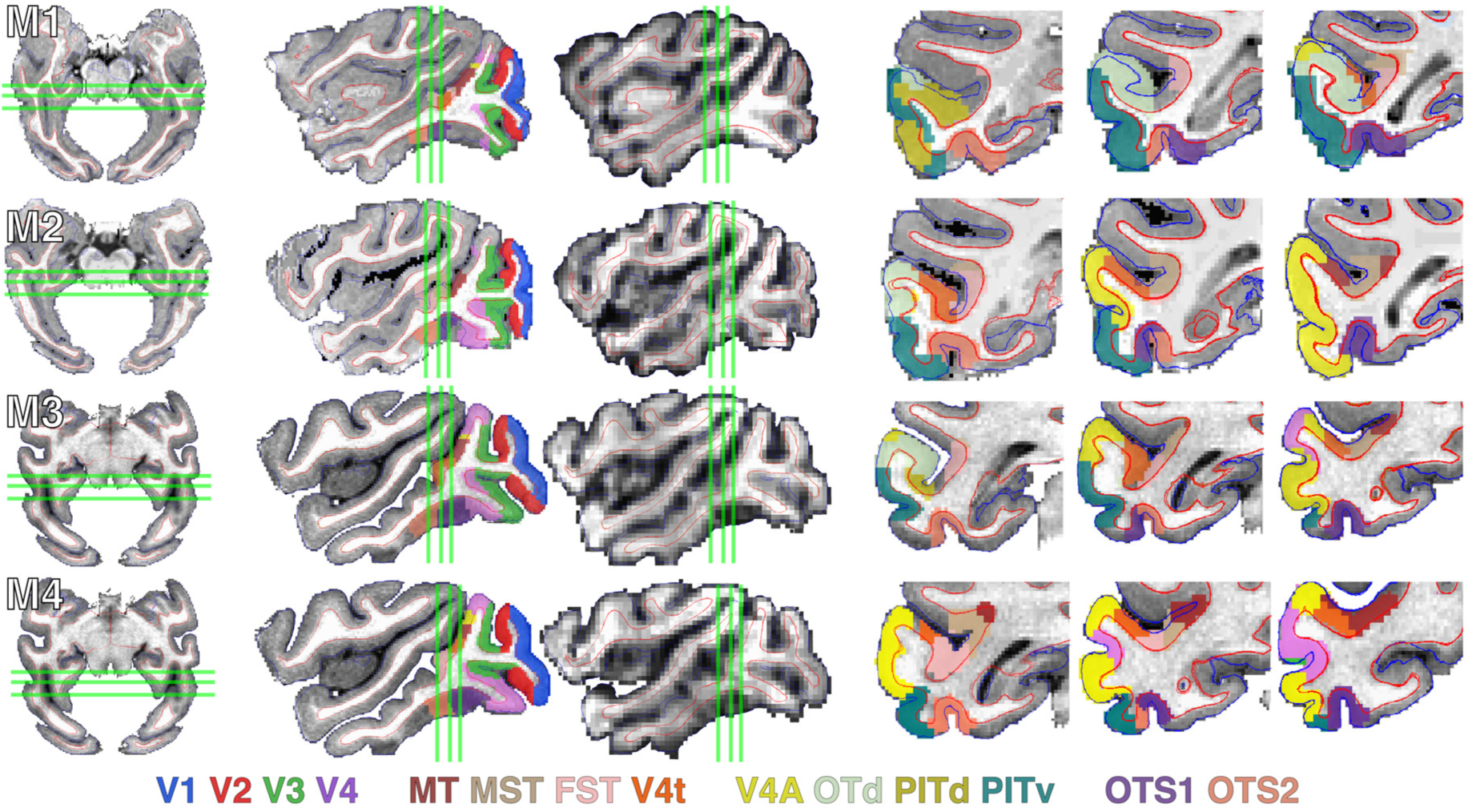
Anatomical localization of retinotopic maps in volume. The location of OTS1 and OTS2 in (left) axial, (middle) sagittal, and (right) coronal views. Mean EPI images from the polar angle experiment are presented aligned to anatomical volumes for the sagittal view. For M1 and M2, EPI images were registered to native anatomical volumes. For M3 and M4, EPI images were registered to the F99 standard template volume. OTS1 and OTS2 as well as V1, V2, V3, V4, V4A, MT, MST, FST, V4t, OTd, PITd, and PITv are illustrated in 3 coronal slices in each monkey. Enlarged views of the left hemisphere of ventral temporal cortex are presented in the coronal slices, evenly spaced at 2mm. Centroids and the shared border of OTS1 and OTS2 fall within these slices, but their area (as well as most other visual maps shown) extend beyond these coronal images. Slices are orientated anterior (left) to posterior (right). Thin blue and red lines in all three views correspond to segmentations of the pial and white matter. Green lines in axial and sagittal slices correspond to the location of coronal slices.

To evaluate the strength of the stimulus-evoked signal relative to noise in OTS1 and OTS2, the response amplitudes were plotted as a function of temporal frequency for the polar angle and eccentricity experiments (Fig 5). For each subject, the response at the stimulus frequency was greater than the responses across all other frequencies, demonstrating a strong link between the measured neural response and stimulus location. For the polar angle experiments, the average percentage signal changes at the SF for OTS1 and OTS2 were 2.25(+/-0.32) and 1.30(+/- 0.32), respectively. For the eccentricity experiments, the average percentage signal changes at the SF for OTS1 and OTS2 were 1.83(+/-0.34) and 1.54(+/-0.27), respectively. For both the polar angle and eccentricity experiments the responses at the stimulus frequency for OTS1 and OTS2 were significantly greater than noise (t(7)>3.74, *p*<0.01). For comparison, the average percentage signal changes for V1, V4, PITd, and PITv, were 5.46(+/-0.43), 7.19(+/-0.65), 2.42(+/-0.43), and 2.83(+/-0.44) from the polar angle experiment and 4.98(+/-0.62), 6.65(+/-0.79), 2.23(+/-0.44), and 3.06(+/-0.45) from the eccentricity experiment. These results demonstrate that visual responses in OTS1 and OTS2 were tightly linked with the stimulus presentation.

**Figure 5.**
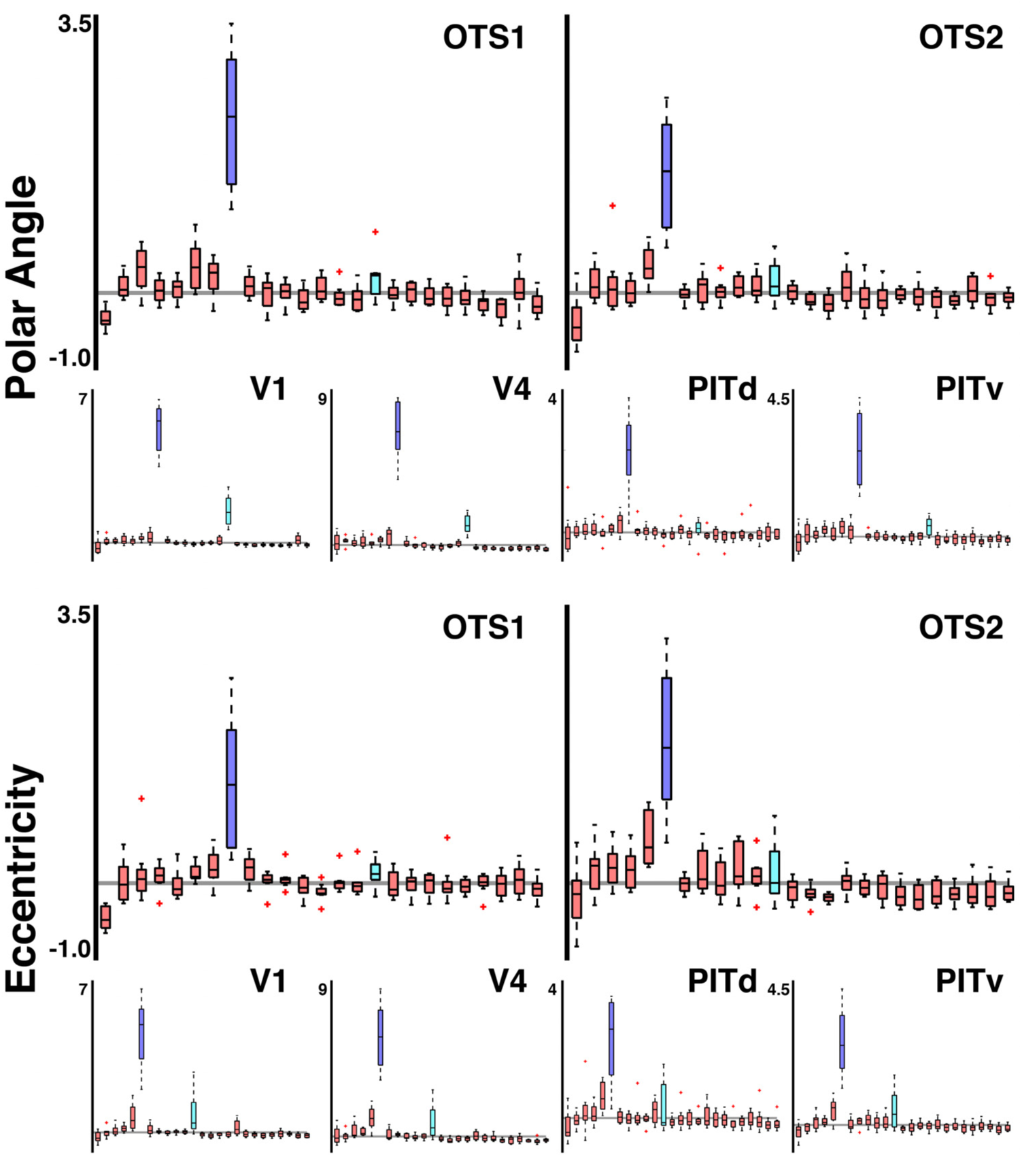
Response amplitude as a function of temporal frequency in OTS1 and OTS2. Box plots of response amplitude converted to % signal change for (top) polar angle and (bottom) eccentricity experiments. The response at the stimulus frequency (dark blue) was significantly greater than the response at all other frequencies. Light blue bars correspond to the first harmonic. For comparison, response amplitudes for areas V1, V4, PITd, and PITv are also shown. Median (black line), interquartiles (whiskers), and 1.5x interquartile outliers (red cross).

OTS1 and OTS2 represented almost exclusively contralateral peripheral space (Fig. 6). 99.0% (+/- 0.01) and 98.0% (+/- 0.01) of voxels preferentially responded to the contralateral visual field in OTS1 and OTS2, respectively (contra vs. ipsi; *p* < 0.0001). 81.0% (+/- 0.05) and 92.4% (+/- 0.04) of voxels responded preferentially to peripheral visual space greater than 6.66° from fixation, and was a significantly larger percentage than representations of foveal (0-3.33°) and parafoveal (3.33-6.66°) in visual space (*p*<0.0001). The disproportionate representation of peripheral space in OTS1 and OTS2 is further apparent in visual comparison to the coverage of areas V1 and V4 (Fig. 6). Note that each data point represents the spatial location of the peak response and does not characterize the spatial extent of responses across the visual field. i.e., the data can be thought of as approximately the center of a voxel’s “receptive field”, not its full visual field coverage. Given that receptive fields span several degrees of visual space in this part of IT(Boussaoud et al., 1991), it is likely that the full visual field is covered by neurons in this area even with this peripheral bias in the spatial distribution of RF centers. A weak preference for the UVF was also apparent in the data. 66.0% (+/- 0.1) and 64.4% (+/- 0.1) of voxels preferentially responded to the UVF, though this bias was not significant (*p* = 0.16 & p = 0.21, respectively). Inspection of individual data revealed that the lack of an effect was due to a single monkey (M1) with all three other monkeys showing a strong preference for the UVF (each monkey > 75%). Due to the limited number of sampled hemispheres (n = 8), more data will be necessary to conclusively evaluate any asymmetry in visual field coverage with respect to UVF and LVF. With the exception of the right hemisphere for M3, all hemispheres for both OTS1 and OTS2 contained representations of both the UVF and LVF, further indicating that both areas should be considered hemifield maps.

**Figure 6.**
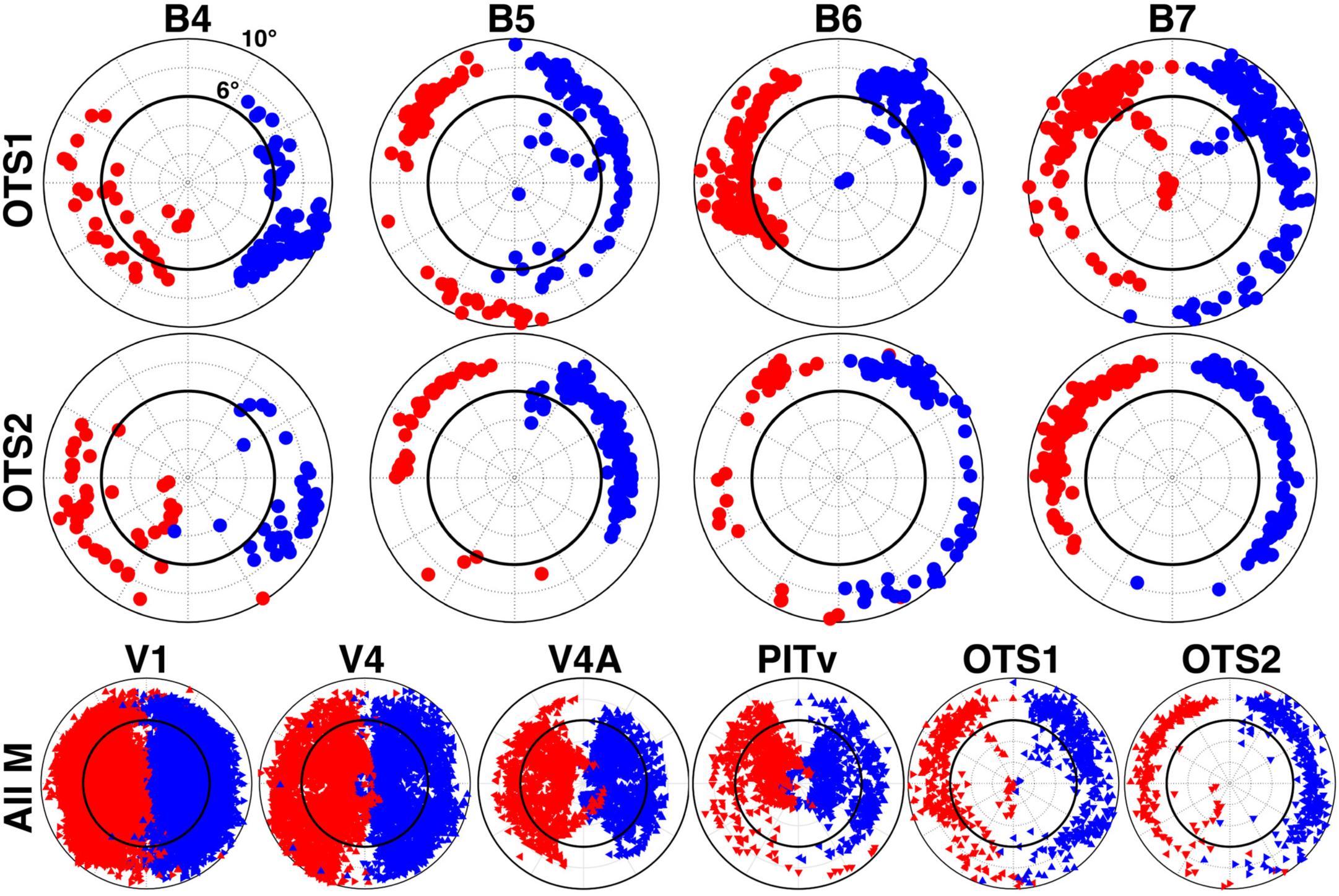
Visual field coverage of OTS1 and OTS2. Individual monkey and group scatter plots of the central 10° visual field representation based on polar angle and eccentricity maps threshold at p < 0.0001. Each point represents the preferred visual field location of a voxel that had significant responses in both polar angle and eccentricity. Red and blue points indicate data from the RH and LH, respectively. Inner solid black line denotes 6° along the eccentricity axis. OTS1 and 2 show a strong contralateral and peripheral preference. Composite visual field coverage is plotted for OTS1 and 2 as well as for areas V1, V4, V4A, and PITv for comparison. For composite data, triangles are presented angled at one of four cardinal orientations for each monkey.

### To split or not to split

OTS1 and OTS2 appear to be distinct areas from surrounding cortex. The criteria for defining a cortical area varies across studies, but typically is based on one or more measures including visual field map organization, chemoarchitecture, anatomical connectivity, and functional response properties(Van Essen, 1985; Gattass et al., 2005). Here, we defined two new areas based on visual field organization and (in a later section) functional response properties. Visual field organization of higher-order visual areas is typically more scattered and contains discontinuities(Gattass et al., 1988; Rosa, 2002), which can make it difficult to resolve areal distinctions. Given the close anatomical proximity to areas V4/V4A/PITv and lack of a clear foveal representation, OTS1 and OTS2 could be interpreted as peripheral extensions of these maps. However, based on the visual field organization of these areas, we think this unlikely. Here, we consider and argue against two alternative accounts (Figure 7).

Given that the OTS has traditionally been considered part of ventral V4 and area TEO/VOT(Lewis and Van Essen, 2000; Gattass et al., 2005), it could be that OTS1 is an extension of ventral V4 and OTS2 is an extension of V4A (Fig. 7, middle). This is unlikely to be the case. The anatomical location and visual field organization of V4 and V4A differs markedly from OTS1 and OTS2, respectively. V4 (and V4A) was located posterior to OTS1 (and OTS2) in all subjects. The lower and upper visual field representations of V4 and V4A are noncontiguous, anatomically separated into dorsal and ventral parts of occipital cortex, respectively. In contrast, the lower and upper visual field representations of both OTS maps are contiguous and there is no dorsal counterpart. As such, the borders of ventral V4 and V4A do not match the borders of OTS1/2. The posterior border of V4 comprises an upper vertical meridian as compared with the lower vertical meridian that forms the posterior border of OTS1. The border between V4 and V4A comprises a HM as compared with the upper vertical meridian that forms the border between OTS1 and OTS2. The anterior border of V4A comprises an upper vertical meridian in contrast to the LVF representation of OTS2. Further, linking OTS1/2 with ventral V4 and V4A would yield quarter-field representations for the central 0-6° of visual field and hemifield representations from 6-10°. The dorsal portions of V4 and V4A fill in the corresponding quarter-fields within 0-6° (Figure 6). Such an areal organization where the UVF and LVF representations are anatomically separated in the central 0-6° **but not in the periphery**, has never been observed and is unlikely to emerge in development(Rosa, 2002). Therefore, V4 and V4A are unlikely to include OTS1 and OTS2.

**Figure 7.**
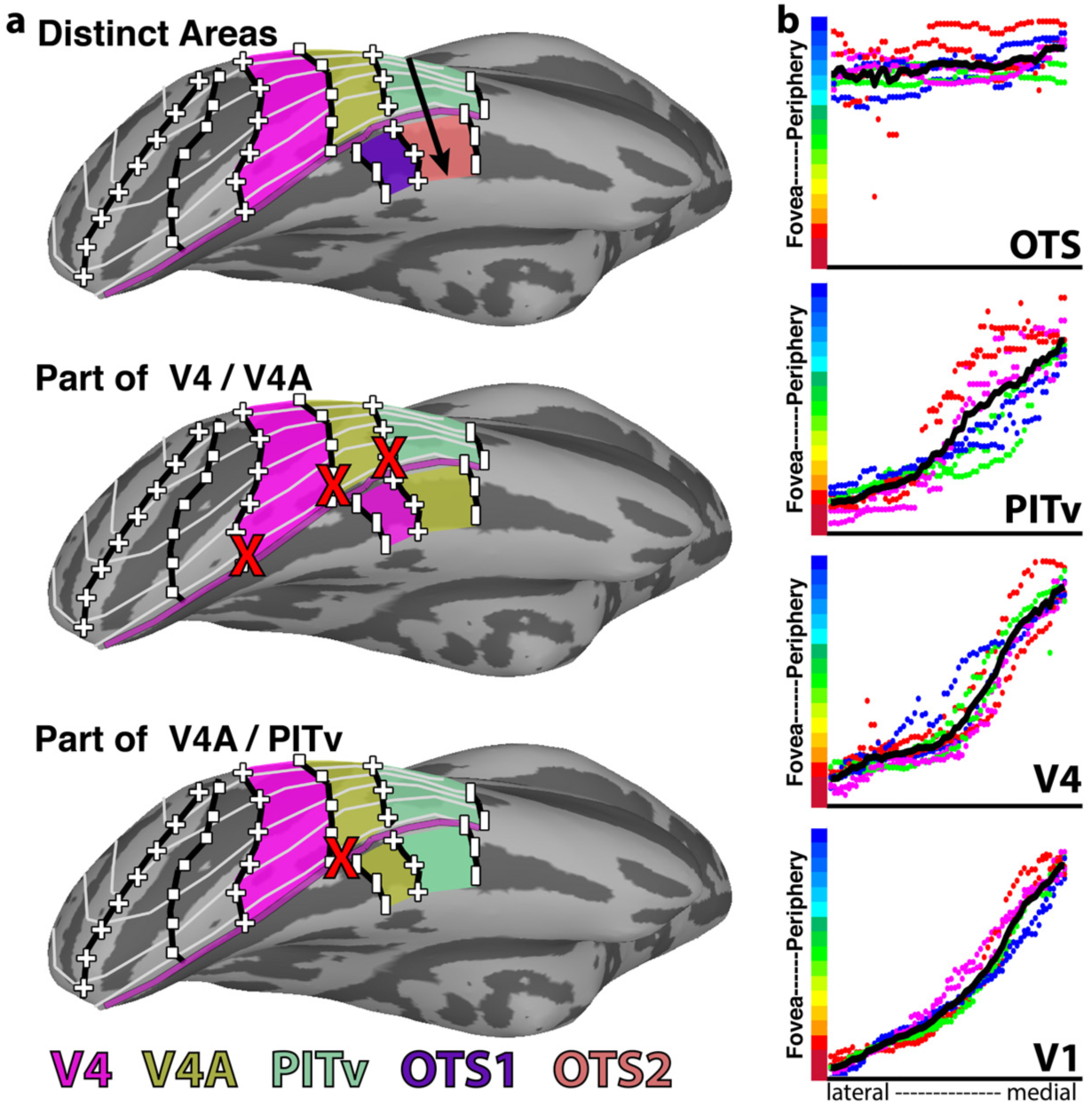
*Alternative models of ventral temporal retinotopic organization.* (a, top) Our proposed model of the retinotopic organization of macaque ventral temporal cortex includes distinct visual field maps for OTS1/2. Black arrow indicates axis of eccentricity map. (a, middle) An alternative model where OTS1 is part of V4 and OTS2 is part of V4A. (a, bottom) A second alternative model where OTS1 is part of V4A and OTS2 is part of PITv. Red x’s mark inconsistencies in polar angle representations between V4/V4A and the OTS maps. Light gray lines illustrate iso-eccentricity lines. The purple line marks the peripheral-most representations for V1-V4, V4A, and PITv. See Figure 1 for other conventions. (b) Eccentricity phase plots from lateral to medial cortex for areas OTS(1 and 2), PITv, V4, and V1. Eccentricity representations progressed from foveal space to the periphery for V1, V4, and PITv. Further medial, there was no clear progression of eccentricity within OTS and the eccentricity line remained flat. Colored dots correspond to individual monkeys (hemispheres plotted separately) and black solid lines correspond to the average across monkeys.

Given the anatomical proximity of V4A and PITv to OTS1 and OTS2, respectively, it could instead be that OTS1 is an extension of V4A and OTS2 is an extension of PITv (Fig. 7, bottom). We think this is also unlikely to be the case. As stated in the preceding paragraph, the quarter-field representation of ventral V4A is incompatible with the hemifield representations of OTS1 and OTS2; the posterior border of V4A comprises a representation of the HM whereas OTS1’s posterior border is the lower vertical meridian (Fig. 2). The polar angle organization of PITv, however, is roughly compatible with that of OTS2. Similar to OTS2, PITv contains a contiguous hemifield representation of contralateral visual space. The polar angle progression along the posterior-anterior axis is similar to OTS2, extending from an upper vertical meridian across the HM to a lower vertical meridian. However, in most hemispheres the visual field representation between the medial border of PITv and the lateral border of OTS2 was noncontiguous. In M1-3, HM representations separated the UVF representation of OTS1/2 from the UVF representations of V4A/PITv. In M2 and M4, the anterior borders of OTS2 and PITv, which correspond to LVF representations, were at acute angles to each and did not align. In M1, the border of OTS2 was separated along the cortical surface by several millimeters from PITv. Further, the eccentricity organization of PITv diverged from that of OTS2 (and OTS1). The eccentricity map of PITv progressed from a foveal representation laterally to peripheral representations at the lateral bank of the OTS (Fig. 7a, purple line) similar to areas V1 and V4 (Fig. 7b). If OTS were part of PITv, eccentricity phase values would continue to progress into the periphery. Instead, the eccentricity representations within OTS remained peripheral, but without any clear progression, suggesting that the PITv map stops at the border with OTS (Fig. 7b). If the foveal representation further medial were part of OTS1 and OTS2, then the eccentricity organization of OTS1 and OTS2 constitutes a reversal of V4A’s and PITv’s eccentricity map. Lastly, visual areas tend to decrease in surface area with increasing distance from V1. The combination of PITv (80.9mm^2^ +/-12.8) and OTS2 would yield a total surface area of 126mm, which is larger than neighboring areas OTd (70.4mm^2^ +/- 3.5) and PITd (67.8mm^2^ +/-5.1) and on par with posterior area V4A (139.0mm^2^ +/-14.6). Thus, it is unlikely that PITv and OTS2 comprise a single area as this would violate the trend for areas to decrease in size moving up the hierarchy. Overall, the organization of the visual field maps of V4A and PITv are inconsistent with OTS1 and OTS2. Therefore, the most likely scenario is that OTS1 and OTS2 are distinct visual field maps, separate from the lateral retinotopic areas.

### Functional response properties

Scene-selective activity was observed within and around the OTS. To test for stimulus selectivity, we showed monkeys blocks of scenes, mosaics of faces, mosaics of bodies, or mosaics of objects; we used mosaics instead of single items in order to cover the entire 20 degrees of the screen, to have comparable retinotopic coverage for all image categories. As seen in Figure 8, a region within the OTS showed stronger activity for scenes vs. faces (top, *p* < 0.00001, FDR corrected) and scenes vs. objects (bottom, *p* < 0.0001, FDR corrected). The spatial location of this scene selectivity matches the location of previously described area LPP(Kornblith et al., 2013). Activations to faces were examined only in 2 of 4 monkeys. The other 2 monkeys were raised with minimal exposure to faces and IT activity specifically to face stimuli was abnormal. Activations to objects and bodies were measured in all 4 monkeys. In general, there was good correspondence between the location of scene selectivity and the extent of OTS1 and OTS2. In 3 of 4 monkeys, scene-selective activations were also observed lateral to area LPP within the PMTS, consistent with anatomical location of the previously identified area mPPA(Nasr et al., 2011). This second scene-selective region lay in area PITv.

OTS1 and OTS2 were highly selective for scenes. In both OTS1/2, there was a main effect of image category (F(3,20) > 9.02, *p* < 0.001), no effect of hemisphere (F(1,20) < 0.34, *p* > 0.5), and no interaction (F(3,20) < 0.1, *p* > 0.95). Activations to scene stimuli were stronger than activations to body or object stimuli (Fig. 6B, t(7) > 4.58, p < 0.01). Activations to scene stimuli were also stronger than activations to face stimuli in the two monkeys in which this was tested (t(3) > 5.5, p < 0.05). To further quantify the stimulus category preference and compare with other visual areas across the visual hierarchy, we calculated a d prime sensitivity index for scene/object condition pairs for V1, V4, PITd, PITv, and OTS1/2. OTS1/2 had the largest d prime indices (> 1.06) with intermediate visual area V4 and ventral temporal area PITv having less than half the selectivity (< 0.48). Even when equating visual field coverage with OTS1/2 by restricting the analysis to peripheral (>6°) representations, d prime indices for OTS1/2 remained double that of PITv (0.44), providing additional evidence that OTS is functionally distinct from PITv. Early visual area V1 and dorsal temporal area PITd had values near zero or slightly negative (i.e., weak object preference). The scene index in OTS1/2 was stronger than any other area (t(7) > 2.56, *p* < 0.05). Taken together, these results demonstrate that OTS1/2 correspond to peak scene selectivity in macaque ventral temporal cortex.

**Figure 8.**
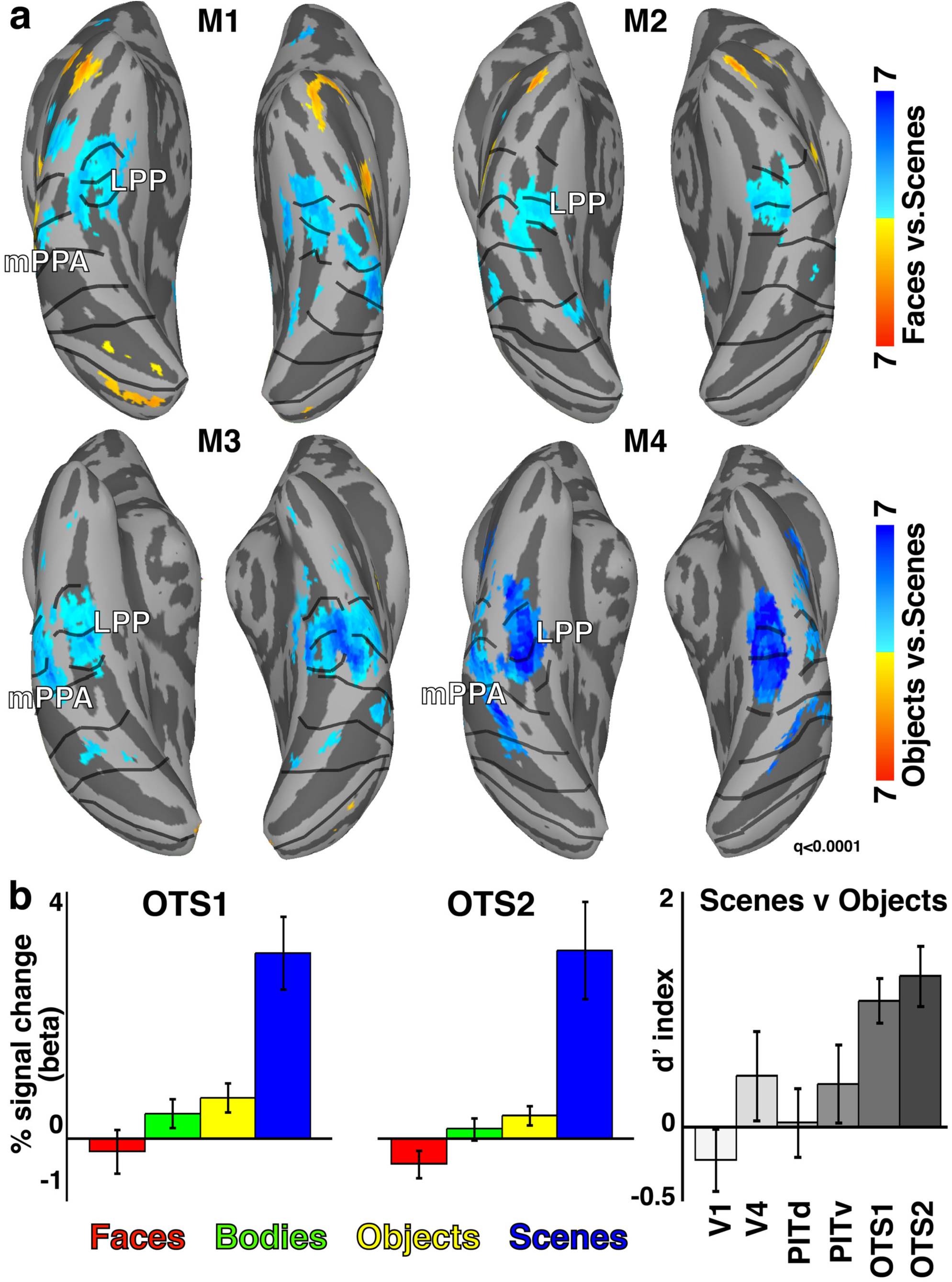
Category selectivity in OTS1 and OTS2. (a) Scene selectivity overlapped with OTS1 and OTS2. This region showed strong selectivity for scenes vs. (top) faces and (bottom) objects. The location of scene selectivity with respect to anatomy and the OTS1 and OTS2 visual field maps was consistent in all four monkeys (p < 0.00001, FDR corrected). (b) Mean responses within OTS1/2 were significantly larger for scene stimuli vs. all other categories (p < 0.01). D primes for distinguishing scenes from objects was significant larger in OTS1/2 than visual areas V1, V4, PITd, and PITv (p< 0.05).

### Retinotopic organization of dorsal scene-selective cortex

Scene-selective responses were also observed in dorsal extrastriate visual cortex. We probed the relationship between scene selectivity and visual field map organization in this region as well. Consistent with prior studies(Fize et al., 2003; Arcaro et al., 2011), we identified several visual field maps within dorsal extrastriate cortex, V3A and DP, and posterior parietal cortex, CIP1/2 and LIP (Fig. 9a). As previously reported with imaging and single unit recordings(Gattass et al., 1988; Arcaro et al., 2011; Arcaro and Kastner, 2015), there is an irregularity in the LVF representation of V3 near areas V3A and DP. The precise organization in this part of dorsal cortex in monkeys is still debated(Lyon and Kaas, 2002; Rosa et al., 2013; Angelucci and Rosa, 2015; Jeffs et al., 2015; Kaas et al., 2015). Further anterior, previous imaging studies reported a single map within the lateral intraparietal sulcus extending along the posterior-anterior axis from an (Fig. 9a, RH #1) upper vertical meridian to a (#2) lower vertical meridian(Patel et al., 2010; Arcaro et al., 2011). Our data suggest that two additional maps exist anterior to the original LIP map, which we have labeled LIP2 and LIP3. The visual field map of LIP2 is a reversal of the polar angle map in LIP1 map, extending from the (#2) lower vertical meridian to (#3) an upper vertical meridian. Anterior to LIP2, we observed weak representations of the HM and another (#4) LVF representation, suggesting the existence of a third map, LIP3. The anterior extent of LIP3 was just posterior to the anterior tip of the IPS (area AIP).

**Figure 9.**
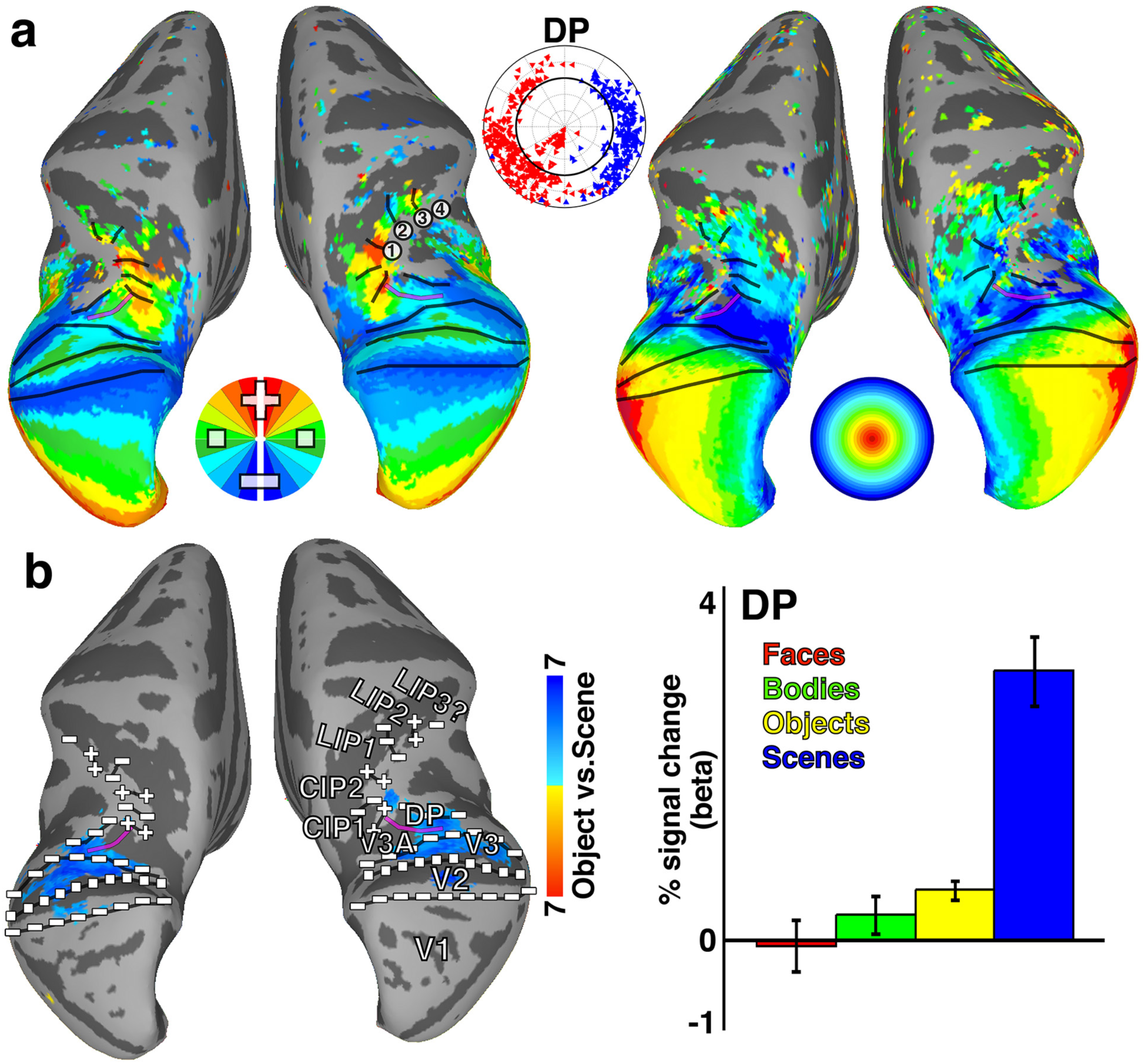
Retinotopic organization of dorsal scene-selective region. (a) Group average (n = 4) polar angle and eccentricity maps of dorsal occipital and posterior parietal cortex. See Figure 1 for conventions. (b) A region within the dorsal prelunate that overlapped with retinotopic areas DP and V3 showed stronger responses to scenes vs. all other categories. This area mainly represented contralateral peripheral space. Our mapping also revealed extensive retinotopic organization throughout the inferior bank of the intraparietal sulcus beyond what has been previously described(Arcaro et al., 2011). Anterior to the map previously referred to as LIPvt (Arcaro et al., 2011)(here referred to as LIP1), we identified another map of contralateral space, referred to as LIP2. Anterior to this map, we saw additional representations of contralateral visual space suggesting the existence of another map, which we tentatively label LIP3.

A focal region of scene-selectivity (vs. objects) in dorsal extrastriate cortex overlapped with visual field maps DP, V3A, as well as part of dorsal V3 (Fig. 9b, left). The location of this dorsal scene selectivity, which Nasr et al.(2011) referred to as mTOS, was consistent with previous studies(Nasr et al., 2011; Kornblith et al., 2013). Visual field map DP responded selectively to scene images and had a category response profile similar to OTS1 and OTS2 (Fig. 9b, right). Similar to the scene-selective maps OTS1/2 in ventral cortex, visual map DP represented peripheral contralateral space (80%, *p* < 0.0001). Notably, in contrast to OTS1/2, DP had a LVF bias (70%, *p* < 0.01). In humans, the dorsal scene area (TOS/OPA) also shows a bias for the lower visual field, and the ventral scene area (PPA), the upper visual field(Silson et al., 2015; Silson et al., 2016). Our data thus further link peripheral visual field representations and scene selectivity, and highlight parallels to the organization in humans.

### Anatomical parcellation of temporal cortex

To further evaluate the consistency of the retinotopic organization across individuals, polar angle and eccentricity data were mapped onto a standard template surface (F99(Van Essen et al., 2012)), and average topographic maps were calculated for each hemisphere (Fig. 10a). Borders between areas were identified at reversals in polar angle representation or eccentricity progression. The group average polar angle and eccentricity maps throughout ventral temporal cortex were remarkably similar to those for individual monkeys, indicating little individual variability in the organization of these visual maps^2^. Group average contrast maps of scene activity vs. object activity (p < 0.0001, FDR corrected in all monkeys) revealed a scene selective region in the occipitotemporal sulcus that overlapped with the group average OTS1 and OTS2 maps (Fig. 10b). These data demonstrate the spatial consistency in the retinotopic organization and category selectivity of OTS across monkeys.

**Figure 10.**
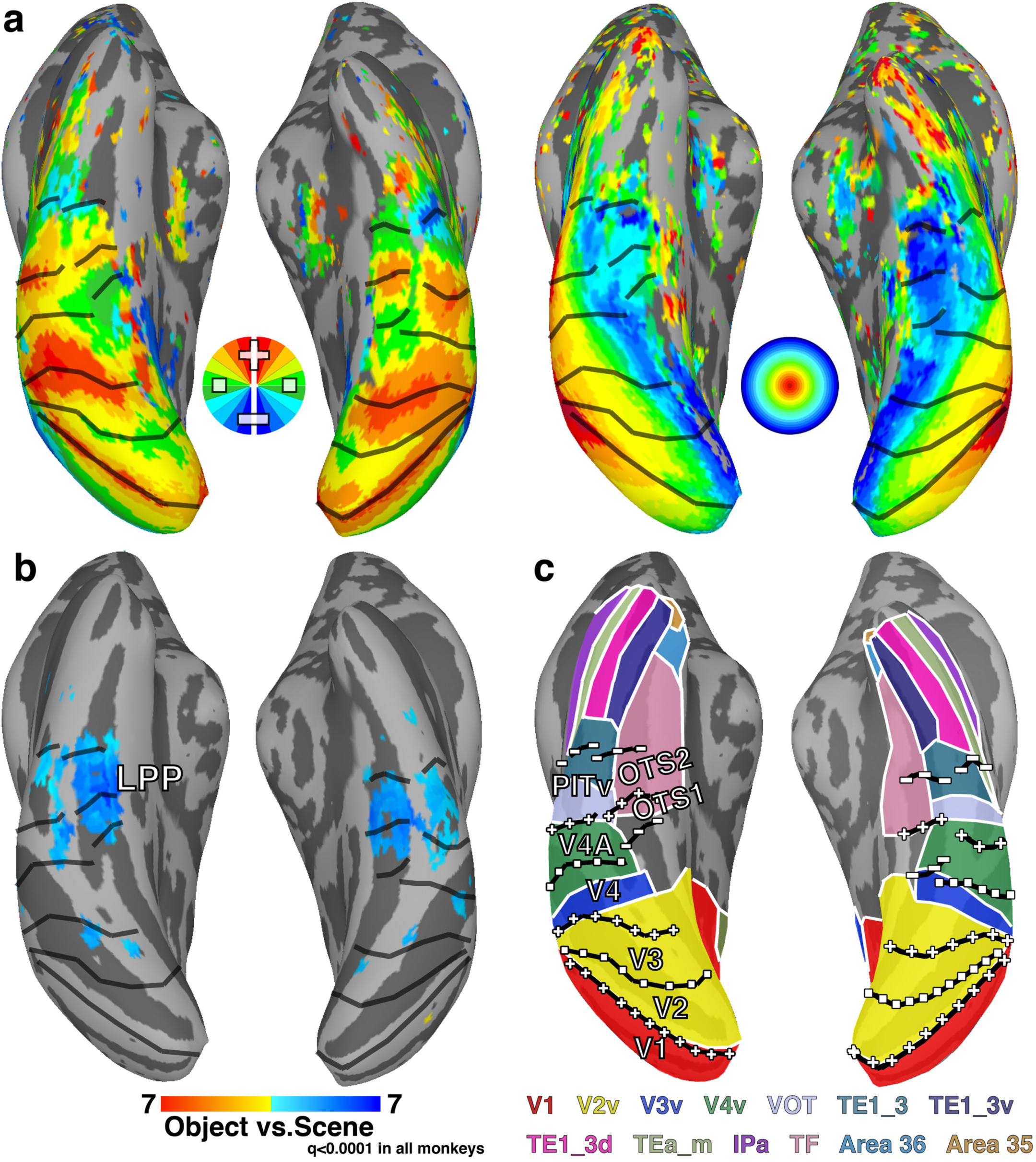
Group average maps. (a) Group average (n = 4) polar angle and eccentricity maps of ventral occipital and temporal cortex. The topographic organization was consistent with individual subject maps indicating that there was good agreement in the representation of visual space across monkeys. Data were threshold at p < 0.001 in each subject. Voxels that were significant in at least 3/4 individuals are displayed. See Figure 1 for conventions. (b) Group average (n =4) map for the contrast of responses to scenes and objects (p<0.0001, FDR corrected). Consistent with the individual subject maps, the group average scene selective region LPP overlapped with OTS1/2. (c) Overlap of retinotopic maps with the Lewis and Van Essen(Lewis and Van Essen, 2000; Van Essen et al., 2012) *atlas. OTS1/2 overlapped with area TF as well as medial portions of V4v and VOT.*

To compare our results with an anatomical and histological parcellation of ventral temporal cortex, area borders from the Lewis and Van Essen atlas(Lewis and Van Essen, 2000; Van Essen et al., 2012) were mapped onto the standard mesh surface and compared to the boundaries determined from the average topography data (Fig. 10c). OTS1 and OTS2 overlapped with the posterior-most portion of area TF as well as medial-most parts of V4. Previous studies have differentiated the posterior portion of area TF, referred to as VTF(Boussaoud et al., 1991), based on visual response properties. OTS1 and OTS2 appear to correspond to area VTF and surrounding cortex. A coarse representation of upper and lower visual space has been previously described in this region of cortex using electrophysiological recordings(Pinon et al., 1998) and anatomical tracers(Sousa et al., 1991), though the orientation of visual field representations along the posterior-axis, the existence of two maps, and their scene selectivity were not described previously. A recent fMRI review(Orban et al., 2014) identified a lower vertical meridian that likely corresponds to the posterior border of OTS1. This review proposed the existence of one retinotopic map posterior to the lower vertical meridian (TFO1) and one anterior (TFO2). The posterior and anterior TFO maps cannot correspond to OTS1 and OTS2, respectively, since the border between the two OTS maps was defined at the upper vertical meridian. It is possible that their anterior TFO map corresponds with OTS1, though only two hemispheres were shown and no additional borders were identified. In this review, a foveal representation was also reported at the lateral extent of this LVF meridian (in one hemisphere, all representations in this area appeared to be foveal). We found no such foveal representation, which, according to their data, should overlap with lateral parts of OTS1/2. This discrepancy between our data is unclear, though it could be due to differences in mapping approaches. In particular, they used phase-encoded mapping stimuli that did not have a break in-between foveal and peripheral-most ring presentations, which can lead to a wrap-around effect in the eccentricity maps.

### Species comparison

The topographic organization and response properties of scene-selective cortex in the macaque paralleled the functional organization in humans. Anterior to human V4 (hV4), four visual field maps have been identified in medial ventral temporal cortex: VO1/2(Brewer et al., 2005) and PHC1/2(Arcaro et al., 2009) (Fig. 11, left). The organization of human PHC, not VO, is similar to macaque OTS. VO1/2 border hV4 and mainly represent the central 8° of visual space visual space. PHC1/2 are located directly anterior to VO1/2 and mainly represent peripheral visual space (> 8°), similar to OTS1/2. Both PHC and OTS maps have larger representations of the UVF (though this did not pass statistical significance tests in OTS), and both progress through the visual field along the posterior-anterior axis starting from a LVF representation, extending to an UVF, then reversing back to the LVF. In humans, the scene-selective region PPA overlaps with PHC1/2 (Fig. 9, right) similar to the overlap between scene-selective LPP and OTS1/2 described above. In both species, these regions are located in major sulci that straddle the lateral extent of parahippocampal cortex. Taken together, these data support a correspondence between monkey OTS1/2 and human PHC1/2. Based on retinotopic organization and category selectivity, there are no other known areas in the monkey that better correspond to PHC in humans and there are no other known areas in the human that better correspond to OTS in monkeys. This proposal of a species homologue between OTS and PHC does not necessarily mean these areas are in 1:1 functional correspondence. While the general functional organization of these areas suggest an evolutionary link, it is entirely possible that novel functional specializations emerged within these regions across the ∼25 million years since these lineages diverged.

**Figure 11.**
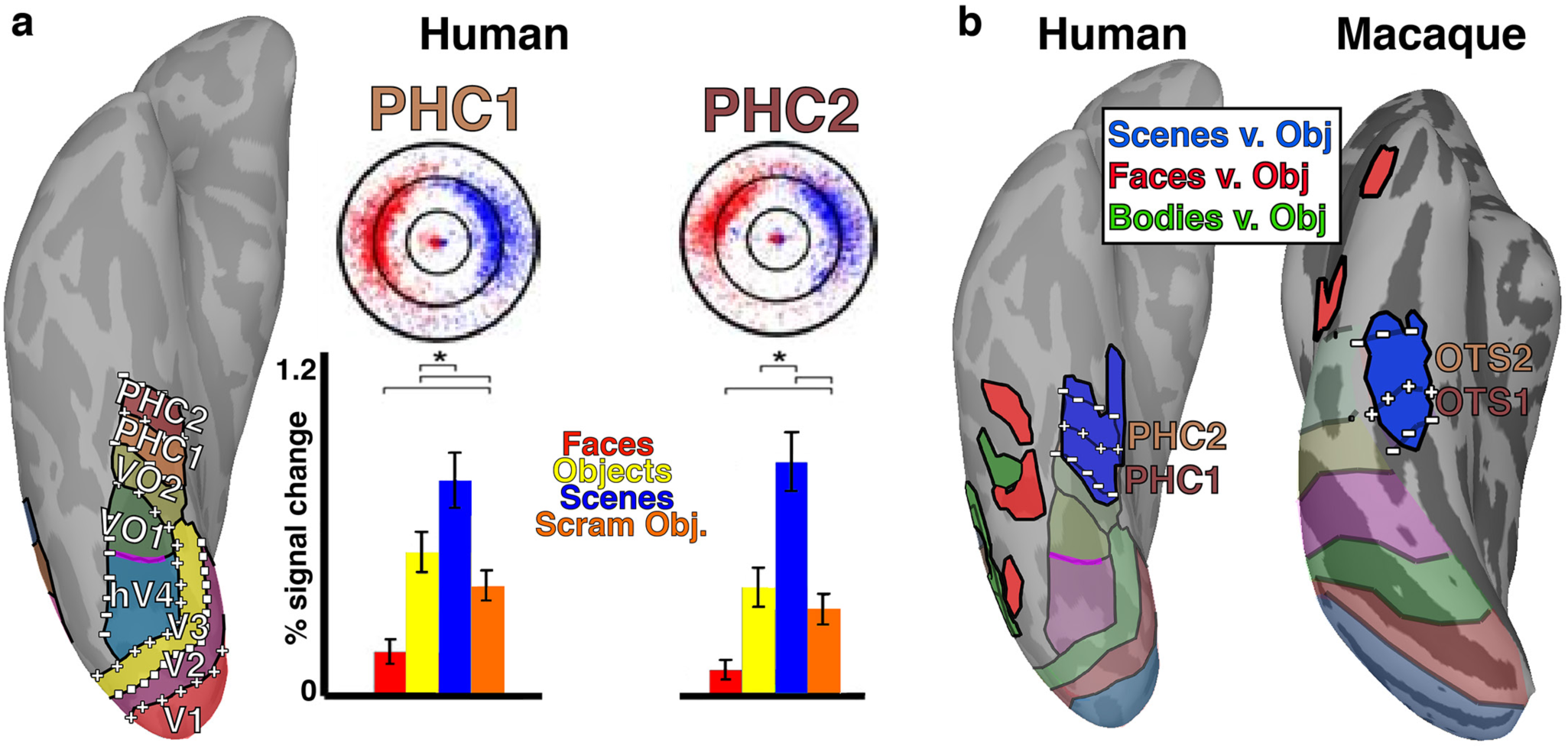
Functional organization of human ventral temporal organization. (a) Visual field maps PHC1/2 in humans have a similar topographic organization to OTS1/2 in monkey. Similar to OTS1/2, PHC1/2 represent peripheral contralateral space and (middle) selectively respond to scene images compared to other image categories. (b) Comparison of ventral temporal functional organization in humans and macaques. A focal region of selectively responsive to scenes vs. objects overlaps with PHC1/2 similar to OTS1/2. Face- and body-selective regions in the human are located lateral to both OTS and PHC in monkeys and humans, respectively. Human data originally published in Arcaro et al. 2009.

The relationship between human VO maps and the monkey visual field maps remains unclear. Human VO directly borders the peripheral representation of hV4. Though the OTS maps were located medial to peripheral representations of V4, it is unlikely that part of OTS also corresponds to VO. The OTS maps were closer to areas V4A and PITv than area V4 (Fig. 2). In some hemispheres, there was a small part of medial ventral temporal cortex that contained representations of contralateral visual space, but did not clearly belong to either the OTS maps or V4/V4A/PITv. It is unlikely that this “gap” between visual field maps corresponds to human VO as this part of cortex mainly represented peripheral space, in contrast to VO’s representation of foveal space(Arcaro et al., 2009). Though V4A is anatomically located in-between V4 and OTS, similar to VO’s position relative to hV4 and PHC in humans, the visual field organization of macaque V4A differs from human VO1/2. As discussed in the **To Split or Not to Split** section, the UVF and LVF representations of monkey V4A are noncontiguous, anatomically separated into dorsal and ventral parts. This organization differs from the contiguous hemifield representation of visual space in human VO1/2. It is possible the visual map organization differs between species as has been proposed for V4(Brewer et al., 2005). It is also possible that PITv corresponds with human VO. PITv contains a contiguous hemifield representation of visual space and a large foveal representation, similar to both VO maps. Further, some scene selectivity is apparent in peripheral portions of the PITv map(Nasr et al., 2011). Similarly, peripheral portions of the VO2 map are selective for scene images(Arcaro et al., 2009). However, monkey PITv is a single map in comparison to the two maps of human VO. It is possible that a single map in macaques has split into two maps in humans. Or it is possible that there is no direct comparison between human VO and the retinotopic organization of temporal cortex in macaques. Given the general correspondence between species for most visual areas, we think this is unlikely and that there is likely a macaque parallel of human VO in functional similarity and relative location in the visual hierarchy even if the topographic organization differs.

The relationship between scene-selectivity and retinotopy in dorsal cortex is also similar between species. In human dorsal visual cortex, scene-selective area TOS/OPA overlaps with several retinotopic areas that largely represent the peripheral LVF(Silson et al., 2015; Wang et al., 2015; Silson et al., 2016). Similarly, our data show that scene-selective representations in dorsal extrastriate cortex (mTOS; (Nasr et al., 2011)) overlapped with several visual field maps that represent peripheral lower visual space. Taken together, these results suggest that the organizations of dorsal and ventral scene-selective regions are preserved across primate species.

## Discussion

We investigated the topographic organization of macaque inferior temporal cortex using standard phase-encoded retinotopic mapping in alert fixating monkeys. By considering both the polar angle and eccentricity organization, we identified two new visual field maps, OTS1 and OTS2, located within the occipital temporal sulcus in medial temporal cortex. These maps were located medial and ventral to previously described visual field maps V4A and PITv(Janssens et al., 2014; Kolster et al., 2014). The polar angle component of the maps was oriented along the posterior-anterior axis and the border between the two maps comprised an upper vertical meridian. OTS1 and OTS2 mainly represented peripheral contralateral visual space and lacked a clear foveal representation, though a patch of central visual field representation further medial may be part of these areas. This retinotopic organization of both areas was consistent in all four monkeys. In addition to the two OTS maps, we identified visual field maps throughout occipital, temporal, parietal and frontal cortex as well as the LGN and pulvinar. Beyond identifying individual visual field maps, polar angle and eccentricity maps covered most of visual cortex. Retinotopically-specific activity almost completely covered visually-evoked activity maps (from the category stimulus experiment) with the exception of anterior temporal STS (AIT), which was visually activated, but did not show clear retinotopic responses. The lack of clear retinotopic response in AIT could reflect a lack of retinotopic organization or methodological limitations (i.e., suboptimal stimulus for mapping large receptive fields with complex selectivity). Altogether, these data demonstrate the prevalence of visual field maps throughout the visual system, including late stages in the ventral visual hierarchy, of macaques.

The retinotopic organization of medial ventral temporal cortex was consistent between humans and macaques. Within and around the OTS, representations of visual space were mainly confined to the periphery with locations of foveal space located laterally within and around the inferior bank of the STS. These data mirror the large-scale eccentricity bias across ventral temporal cortex previously reported in humans(Hasson et al., 2002) and in monkeys(Lafer-Sousa and Conway, 2013). Within this broad eccentricity organization, there were parallels between species in the organization of individual visual maps. The organization of OTS1 and OTS2 corresponds with visual field maps PHC1 and PHC2, respectively, in the human. Both human and monkey maps mainly represent peripheral space and are biased towards the UVF. In humans, PHC1 and PHC2 have been proposed to comprise a visual field map cluster(Arcaro et al., 2009). Typically, visual field map clusters are made up of functionally similar individual maps that share a common eccentricity map, with the borders between areas defined at reversals along the polar angle dimension(Wandell et al., 2005). Given the parallels in organization between OTS and PHC, it is possible that OTS1/2 are also part of a visual field map cluster. However, we did not identify a foveal representation within OTS1/2, which is a defining feature of clusters, though foveal representations were medial to OTS1/2 in most monkeys. Overall, our data demonstrate that the retinotopic organization of macaque ventral medial IT cortex matches well with that in humans by several criteria.

Our results demonstrate a relationship between scene-selective areas in humans(Aguirre et al., 1998; Epstein and Kanwisher, 1998) and primates(Nasr et al., 2011; Kornblith et al., 2013). Within ventral temporal cortex, previous studies have reported three scene-selective regions, mPPA, LPP and MPP, in monkeys(Nasr et al., 2011; Kornblith et al., 2013), but only one ventral region, PPA, in humans(Aguirre et al., 1996; Epstein and Kanwisher, 1998). Though response properties in these monkey regions are similar to responses in the human PPA, it has remained unclear which, if any, of the monkey scene-selective areas is comparable to the human PPA. Here, the retinotopic organization of ventral temporal cortex disambiguates these regions. Visual maps OTS1/2 overlapped with the functionally defined scene area, LPP. This correspondence between visual field organization and category selectivity in monkeys mirrored the overlap between visual maps PHC1/2 and scene-selective area PPA in humans. Our data provide new evidence that monkey LPP is the homologue to human area PPA. There is an additional scene selective region in human (TOS/OPA) and macaque dorsal cortex (mTOS), and these areas in both species are biased towards ventral visual field(Silson et al., 2015; Silson et al., 2016). Taken together, our data suggest the functional organization of scene-selective cortex is preserved across primate species.

Our data suggest that OTS1/2 formed early in development. While retinotopic mapping is typically performed in adults, we scanned juvenile monkeys (∼1.5 yrs.). Already at 1.5 years of age, the retinotopic organization of visual cortex matched that found in the adult. Two of our monkeys were also raised in abnormal conditions where they were restricted from seeing a particular image category. Though this early abnormal experience drastically altered responses to that visual category in IT(Arcaro et al., submitted), the organization of visual field maps within IT were comparable with normally reared monkeys. Further, recent results from our lab suggest that the general retinotopic organization of cortex is present at birth and is likely instrumental in guiding experience-dependent development in IT(Arcaro and Livingstone, in revision). We found that the peripheral retinotopic organization of OTS1/2 is established very early in development, potentially prenatally, and therefore could guide the subsequent clustering of scene selectivity, since scenes are experienced peripherally(Hasson et al., 2002; Torralba and Oliva, 2003). Such a mechanism could explain where on the large-scale eccentricity map in IT(Hasson et al., 2002; Lafer-Sousa and Conway, 2013) domains form(Srihasam et al., 2014) and why the anatomical location of scene selective domains relative to the locations of other function clusters is so consistent across individuals as well as species.

The retinotopic organization of individual visual field maps can also account for where along the anterior-posterior axis category domains form in IT. There is a growing literature on the correspondence between retinotopic maps and category-selective domains. In humans, ventral scene-selective area PPA overlaps with retinotopic maps PHC1 and PHC2 (Arcaro et al., 2009) and dorsal scene-selective area TOS/OPA overlaps with V3B and IPS0(Bettencourt and Xu, 2013; Silson et al., 2016). Lateral object-selective area LOC partially overlaps with LO1 and LO2(Sayres and Grill-Spector, 2008) and lateral occipital face-selective area OFA partially overlaps with retinotopic areas PITd and PITv(Janssens et al., 2014). In monkeys, the posterior (PL) and medial (ML) face patches partially overlap retinotopic maps OTd and PITd(Janssens et al., 2014). Though this prior study reported that ML was differentiable from PITd, we find much more of an overlap between the two. Here, we show that ventral scene-selective area LPP overlaps with OTS1 and OTS2 and dorsal scene-selective area mTOS overlaps with areas DP, V3A, and V3d. Across all of these studies, domains overlapped with 2 or more retinotopic maps that were consistent across subjects. Therefore, even though there appears to be no 1:1 correspondence between individual category domains and retinotopic maps, the organization of visual field maps is still predictive of the location of individual category domains. This correspondence likely has functional relevance. While the large-scale eccentricity organization in primate IT guides where along the lateral-medial axis category domains are localized, we propose that the relation to individual visual field maps anchor these domains along the anterior-posterior axis, effectively reflecting where along the visual hierarchy these domains emerge. Such a mechanism can explain the localization of novel domain formation in IT from intensive training(Srihasam et al., 2014). In this study, the location of novel domains varied along the lateral-medial axis, reflecting differences in eccentricity and low-level stimulus properties such as curvature and spatial frequency. However, the anterior-posterior locations were similar across domains and with naturally occurring domains (e.g., face and scene domains). It is likely that at this level of the visual processing hierarchy, the conditions are ripe for clustering of information along category boundaries or visual features that systematically vary across categories. Thus, the combination of a large-scale eccentricity map and individual visual field maps distributed along the anterior-posterior extent guide the stereotyped localization of category domains in IT. Altogether, retinotopy is clearly a fundamental organizing principle of the ventral stream.

## Acknowledgments

This work was supported by NIH grants RO1 EY 25670, P30 EY 12196, and F32 EY 24187. This research was carried out in part at the Athinoula A. Martinos Center for Biomedical Imaging at the Massachusetts General Hospital, using resources provided by the Center for Functional Neuroimaging Technologies, P41EB015896, a P41 Biotechnology Resource Grant supported by the National Institute of Biomedical Imaging and Bioengineering (NIBIB), National Institutes of Health, and NIH Shared Instrumentation Grant S10RR021110. We thank A. Schapiro for helpful comments on the manuscript. We thank P. Schade for monkey training and for help in scanning.

## Footnotes

1 The lack of a clear vertical meridian representation in a few hemispheres likely reflects an fMRI sampling bias due to the small cortical extent, and should not be taken as evidence for lack of a vertical meridian representation. Voxels that sample from neurons with RFs centered on the vertical meridian also sample neurons with RFs positioned off the vertical meridian in neighboring parts of contralateral visual space, resulting in a representation at the voxel level off the vertical meridian. This effect increases moving up the visual hierarchy, as RFs get larger and the cortical extent of areas decrease.

2 There was a general tendency for the vertical meridians to be under-represented, especially for anterior visual maps. Vertical meridian representations were apparent in individual monkeys (Fig. 2), thus this is likely an artifact due to averaging. As a result, the visual field map of areas with hemifield representations will compress towards the HM (i.e., have an under representation of the vertical meridians). The visual field map of areas with a quarter-field representation will compress towards the midpoint of the HM and VM in that quadrant. This effect increases moving up the visual hierarchy as areal size (and the extent representing meridians) decreases.

